# Proximity proteome mapping reveals PD-L1-dependent pathways disrupted by anti-PD-L1 antibody specifically in EGFR-mutant lung cancer cells

**DOI:** 10.1101/2021.09.17.460833

**Authors:** Anudari Letian, Eyoel Yemanaberhan Lemma, Paola Cavaliere, Noah Dephoure, Nasser K. Altorki, Timothy E. McGraw

## Abstract

PD-L1, a transmembrane ligand for immune checkpoint receptor PD1, has been successfully targeted to activate an anti-tumor immune response in a variety of solid tumors, including non-small cell lung cancer (NSCLC). Despite the success of targeting PD-L1, only about 20% of patients achieve a durable response. The reasons for the heterogeneity in response are not understood, although some molecular subtypes (e.g., mutant EGF receptor tumors) are generally poor responders. Although PD-L1 is best characterized as a transmembrane PD1 ligand, the emerging view is that PD-L1 has functions independent of activating PD1 signaling. It is not known whether these cell-intrinsic functions of PD-L1 are shared among non-transformed and transformed cells, if they vary among cancer molecular subtypes, or if they are impacted by anti-PD-L1 therapy. Here we use quantitative microscopy techniques and APEX2 proximity mapping to describe the behavior of PD-L1 and to identify PD-L1’s proximal proteome in human lung epithelial cells. Our data reveal growth factor control of PD-L1 recycling as a mechanism for acute and reversible regulation of PD-L1 density on the plasma membrane. In addition, we describe novel PD-L1 biology restricted to mutant EGFR cells. Anti-PD-L1 antibody treatment of mutant EGFR cells perturbs cell intrinsic PD-L1 functions, leading to reduced cell migration, increased half-life of EGFR and increased extracellular vesicle biogenesis, whereas anti-PD-L1 antibody does not induce these changes in wild type EGFR cells. These specific effects in mutant EGFR cells might contribute to the poor anti-PD-L1 response of mutant EGFR tumors.

## Introduction

Therapies targeting the PD1 / PD-L1 checkpoint axis are effective in many solid tumors, including non-small-cell lung carcinoma (**NSCLC**)^1,2^. Despite the successes of these therapies, only about 20% of patients achieve durable response. The reasons for the heterogeneity in response are not understood. Cancer cell PD-L1 expression has proven to be an imperfect biomarker of response to immune checkpoint therapy^3^. Some molecular subtypes, most notably mutant EGFR NSCLC but ALK rearrangement tumors as well^4–6^, respond poorly to anti-PD-L1 antibody therapy although the reason(s) for the resistance have not been described.

Although PD-L1 is best characterized as a transmembrane PD1 ligand, there is accumulating evidence that PD-L1 has functions independent of activating PD1 signaling. Recent studies support a role for PD-L1 in regulation of epithelial mesenchymal transition (**EMT**), resistance to chemotherapeutic agents as well as interferon-mediated cell toxicity^7–10^. It is not known whether these cell-intrinsic functions of PD-L1 are shared among non-transformed and transformed cells, if they vary among cancer molecular subtypes, or if they are impacted by anti-PD-L1 therapy.

In addition to being localized to the plasma membrane, PD-L1 will transit through a number of membrane compartments during its life time. CMTM6 has been described as a protein that contributes to the control of PD-L1 expression by regulating PD-L1 targeting to lysosomes^11,12^. Additional proteins such as HIP1R and TRAPCC4 complex have been shown to target PD-L1 for lysosomal degradation or Rab11-mediated recycling to the plasma membrane respectively^13,14^. ALIX, an ESCRT accessory protein, has been shown to sort PD-L1 into intralumenal vesicles of MVBs and control its abundance on extracellular vesicles^15^. Although these studies show PD-L1 degradation or trafficking can be modified through the expression or depletion of these proteins, it remains unclear whether steps of PD-L1 trafficking are regulated under physiologic conditions and how physiologic control of PD-L1 trafficking contributes to its function as an immune checkpoint or to its cell-intrinsic activities.

Here we use quantitative microscopy techniques and APEX2 proximity mapping to describe the behavior of PD-L1 in living human lung epithelial cells. APEX2 biotinylation mapping is a method to define the proteome within 20nm of a protein linked to the APEX peroxidase^16,17^. We leveraged the PD-L1 proximity map to provide a context for further investigations of potential cell-intrinsic roles of PD-L1. Our data reveal growth factor control of PD-L1 recycling as a mechanism for acute and reversible regulation of PD-L1 density on the plasma membrane. In addition, we describe novel PD-L1 biology restricted to mutant EGFR cells. Anti-PD-L1 antibody treatment of mutant EGFR cells perturbs cell intrinsic PD-L1 functions, leading to reduced cell migration, increased half-life of EGFR and increased extracellular vesicle biogenesis, whereas anti-PD-L1 antibody does not induce these changes in wild type EGFR cells.

## Results

### PD-L1 is dynamically maintained on the plasma membrane of cells

PD-L1 was distributed between the plasma membrane (**PM**) (immunostaining of non-permeabilized cells) and intracellular compartments (immunostaining of permeabilized cells) of freshly isolated primary human NSCLC cells (Fig.1A). Of the total cell PD-L1, approximately 30% was at the PM (Fig.1B). The steady-state PD-L1 distribution between PM and interior of primary NSCLC cells was similar to its distribution in established NSCLC cell lines as well as its distribution in a non-transformed human lung epithelial cell line (BEAS-2B) (Fig.1C). Because oncogenic KRAS and EGFR are the two most common mutations in NSCLC^18^, we focused our studies on cell lines harboring either KRAS or EGFR mutations. PD-L1’s distribution did not vary with the oncogenic driver, nor was it influenced by total expression of PD-L1 (Fig.1B-D). In primary NSCLC cells and all the NSCLC cell lines examined, PM PD-L1 showed some enrichment in puncta, which was enhanced by bivalent secondary antibody staining compared to monovalent secondary Fab stain (Fig.S1A). Intracellular PD-L1 was primarily in vesicles dispersed throughout the cytoplasm, although in some cells there was a juxta-nuclear concentration typical of the Trans-Golgi Network/Golgi (Fig.1A, E, Fig.S1B). In all NSCLC cell lines examined, interferon gamma (IFNγ), a potent transcriptional regulator of PD-L1^19,20^, increased the total amount of PD-L1 protein without affecting PD-L1 distribution between the interior and PM of cells, its localization to PM puncta, or its localization to cytoplasmic vesicles (Fig.1E-F, Fig.S1B).

**Figure 1.**
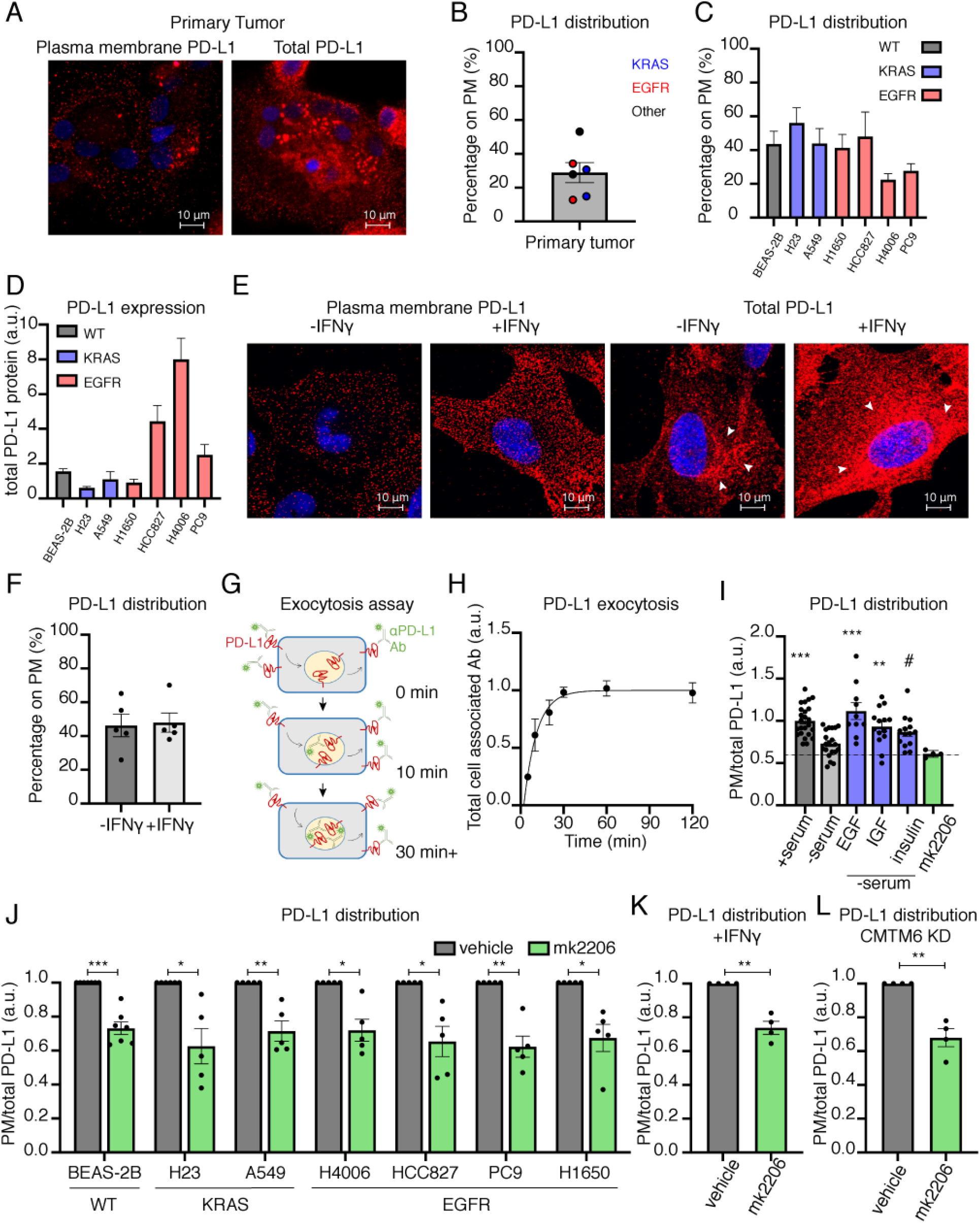
PD-L1 is dynamically maintained on the plasma membrane of cells. (A) Representative confocal images of primary human tumor cells immunostained for plasma membrane (PM) and total PD-L1. Red: PD-L1. Blue: Hoechst. (B) Quantification of percentage of PM PD-L1 in primary human tumors. Each data point is a patient sample color coded for known oncogenes. (C-D) Quantification of (C) percentage of PM PD-L1 and (D) total PD-L1 protein expression in human lung cell lines. Cell lines in blue and red harbor mutant KRAS and mutant EGFR respectively. (E) Representative confocal images of BEAS-2B with or without IFNγ treatment and immunostained for PM and total PD-L1. Red: PD-L1. Blue: Hoechst. Sum intensity of z-stacks is projected. Fluorescence intensity is equally scaled across panels. (F) Quantification of PM localization of PD-L1 in BEAS-2B cells treated with IFNγ. Unpaired student’s t-test. (G) Schematic of PD-L1 exocytosis assay. Live cells are incubated with unlabeled PD-L1 antibody. Cells are fixed, permeabilized and stained with fluorescently labeled secondary antibody depicted as as a star. (H) PD-L1 steady state exocytosis in BEAS-2B cells. The line is a fit of the data to one-phase association exponential equation, N=3. (I) PD-L1 PM to total ratio with growth factors treatment or Akt inhibition in BEAS-2B cells. One-way ANOVA and post-hoc Tukey test comparing each condition to -serum. (J) PD-L1 PM to total ratio in cell lines treated with vehicle (DMSO) or Akt inhibitor for 4 hrs. One sample t-test. (K) PD-L1 PM to total ratio in IFNγ treated BEAS-2B cells incubated with vehicle (DMSO) or Akt inhibitor. One sample t-test. (L) PD-L1 PM to total ratio in CMTM6 knocked-down (KD) IFNγ-treated BEAS-2B cells with vehicle (DMSO) or Akt inhibitor. Except panel B and H, each data point is an experiment. All error bars = SEM. * ≤ 0.05, ** ≤ 0.01, *** ≤0.001, # = 0.058

PM PD-L1 is poised to function as an immune checkpoint by binding in trans to PD1 on the surface of T cells^21,22^. Intracellular PD-L1 could serve as a reservoir to rapidly modulate the amount of PD-L1 on the PM to fine-tune its function as an immune checkpoint on a shorter timescale than IFNγ transcriptional regulation of PD-L1 expression. To interrogate the relationship between the PM and intracellular pools of PD-L1, we used a quantitative fluorescence microscopy assay in which trafficking of PD-L1 between intracellular compartments and the PM is monitored by incubating living cells in medium containing an antibody that binds to the extracellular domain of PD-L1 (Fig.1G). At early time points of incubation, the amount of cell-associated anti-PD-L1 antibody predominantly reflects the amount of PD-L1 on the PM. An increase in cell-associated anti-PD-L1 antibody with increasing time of incubation reflects antibody binding to unbound PD-L1 newly recycled to the PM from intracellular compartments. Cell-associated anti-PD-L1 antibody will increase until all the intracellular PD-L1 that is in equilibrium with the PM has cycled to the PM (i.e., bound by anti-PD-L1 antibody). The rise to the plateau is the rate at which intracellular PD-L1 traffics to the PM. Cell-associated anti-PD-L1 antibody reached a plateau by 30 min, demonstrating that PM density of PD-L1 is dynamically maintained by constitutive PD-L1 cycling between intracellular compartments and the PM. The equilibration halftime for PD-L1 exchange between the PM and intracellular compartments is approximately 10 min (Fig.1H). These data suggest PD-L1’s traffic to and from the PM as a mechanism to acutely and reversibly increase or decrease PM PD-L1 amounts, an effect that would impact PD-L1 immune checkpoint function.

We tested the hypothesis that PD-L1 trafficking is acutely regulated by determining the effect of growth factor stimulus on PM PD-L1. Removal of tonic growth factor signaling by incubation of cells in serum-free medium for 4 hrs resulted in a significant reduction of PM PD-L1 (Fig.1I). This reduction of PM PD-L1 reflected a redistribution of PD-L1 from the PM to intracellular compartments without significant degradation of PD-L1 (Fig.S1C). Addition of different growth factors (i.e., EGF, IGF or insulin) to serum-free medium restored PM PD-L1 without affecting the total amount of PD-L1 (Fig.1I, Fig.S1C). Thus, signaling downstream of various growth factors actively maintains PD-L1 on the PM by regulation of PD-L1 cycling to and from the PM, and identifying acute modulation of growth factor signaling as a mechanism that contributes to the control of PD-L1 PM. This acute and reversible effect of growth factors stimulation reflects a change in PD-L1 localization, not a change in total PD-L1 amounts.

Growth factors commonly regulate membrane protein trafficking downstream of activation of the serine/threonine AKT kinase^23,24^. Consistent with a role for AKT in actively maintaining PD-L1 on the PM, inhibition of AKT reduced the amount of PD-L1 in the PM without affecting the total amount of PD-L1 expressed in cells (Fig.1I, Fig.S1C-E). Inhibition of AKT resulted in a near two-fold reduction of the PD-L1 recycling rate constant, accounting for reduced PM PD-L1 (Fig.S1D). The effect of AKT inhibition on PD-L1 expression in the PM was independent of the driver oncogene (Fig.1J). AKT-dependent regulation of PM PD-L1 was distinct from IFNγ transcriptional regulation of PD-L1 expression; therefore, it is a complementary mode to regulate PM density of PD-L1 (Fig.1K).

CMTM6, a ubiquitously expressed protein, post-translationally controls PD-L1 PM amounts by determining whether internalized PD-L1 is recycled or targeted for degradation^11,12^. In agreement with those studies, transient knockdown of CMTM6 resulted in a 30% reduction of total PD-L1 amounts. However, the distribution of the remaining PD-L1 between the plasma membrane and interior of cells was unchanged (Fig.S1F). Regulation of PM PD-L1 amounts downstream of AKT was also unaffected by CMTM6 knocked down. Thus, regulation of PD-L1 traffic downstream of AKT is a mechanism to acutely modulate PM PD-L1 independently of other known mechanisms (e.g., IFNγ and CMTM6) (Fig.1L).

### PD-L1 proximity mapping

Localization of PD-L1 to intracellular compartments might reflect cell-intrinsic function(s) of PD-L1 that are in addition to regulating the fraction of PD-L1 expressed on the surface of cells. A more complete knowledge of the proteins in proximity to PD-L1 will provide a framework for understanding potential cell intrinsic functions of PD-L1. To identify the PD-L1 proximal proteome we used APEX2 mapping^17^. We expressed a PD-L1 chimera in which the APEX2 peroxidase catalytic domain was fused to the cytosolic carboxyl terminus of PD-L1 (PD-L1-APEX2) (Fig.2A). PD-L1-APEX2 expressed in BEAS-2B cells behaved similarly to endogenous PD-L1. Approximately 40% of PD-L1-APEX2 was in the PM, localized to puncta, whereas intracellular PD-L1-APEX2 was distributed between dispersed puncta and a concentration in the peri-nuclear region of cells (Fig.2B-C) Furthermore, the amount of PD-L1-APEX2 in the PM was controlled by AKT (Fig.2D). These data validate PD-L1-APEX2 as a reporter for PD-L1.

**Figure 2.**
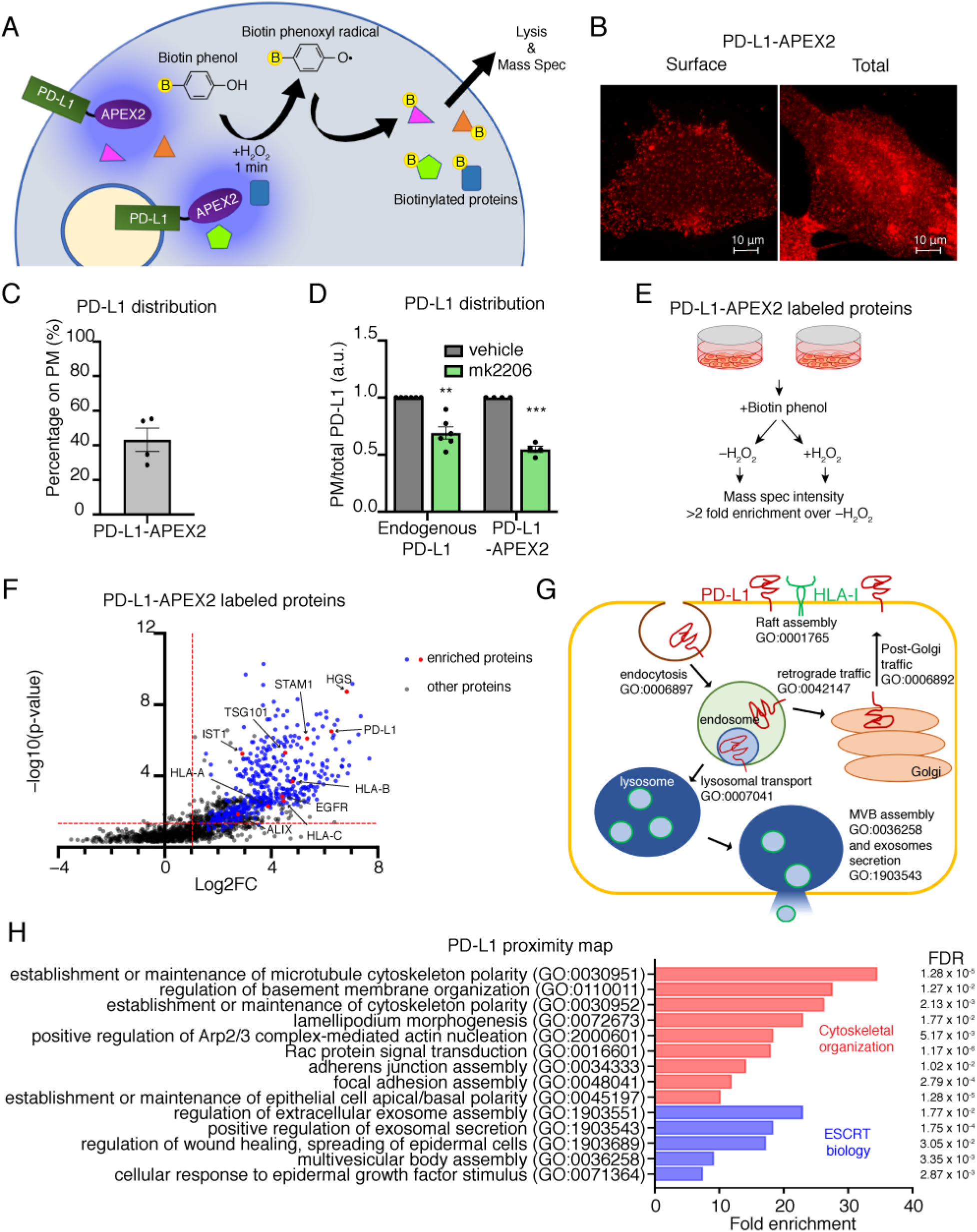
PD-L1 proximity mapping. (A) Schematic of PD-L1-APEX2 labeling assay. (B) Representative epifluorescence image of PD-L1-APEX2 plasma membrane and total stain in BEAS-2B cells. (C) PM to total distribution of PD-L1-APEX2 in BEAS-2B cells. (D) PM to total ratio of endogenous PD-L1 and PD-L1-APEX2 construct in BEAS-2B cells treated with vehicle (DMSO) or Akt inhibitor. One-sample t -test. (E) Schematic of experimental design for PD-L1-APEX2 proximity labeling. (F) Volcano plot of PD-L1-APEX2 labeled proteins. FC = fold change of (labeled/–ctrl) spectral intensity. Dashed red lines represent 4-fold change on the x-axis, and p-value corresponding to FDR=0.1 after Benjamini-Hochberg correction. (G) Biological pathways and compartments identified in PD-L1-APEX2 proximity labeling. David gene ontology terms and identifiers are shown. (H) Additional Biological processes pathways enriched in PD-L1-APEX2 labeling in BEAS-2B. All error bars = SEM. * ≤ 0.05, ** ≤ 0.01, *** ≤0.001.

To catalog the PD-L1 proximal proteome, living cells were pre-incubated with biotin phenol, and the reaction was catalyzed by the addition of hydrogen peroxide to generate a biotin-phenoxyl radical that covalently binds to electron rich amino acids on nearby proteins (Fig.2E, Fig.S2A). PD-L1 proximal proteins were identified by streptavidin beads enrichment and mass spectrometry analysis (Fig.2E). In each of the five individual experiments, APEX2 biotinylated proteins were identified as streptavidin-absorbed proteins whose intensities in the H_2_O_2_-treated samples were 2 fold or greater than in samples not treated with H_2_O_2_ (no APEX reaction). The aggregate PD-L1 proximity proteome was determined as the overlapping set of 438 proteins from the five independent experiments (Fig.2F, Table S1). Of note, PD-L1 (CD274) was identified as an APEX2 biotinylated protein in all five experiments.

HLA class I proteins (HLA-A, -B and -C), which are involved in presentation of endogenous antigens, were identified as components of the PD-L1 proximity proteome (Fig.2F,G, Table S1). Immunofluorescence confirmed the proximity of PD-L1 to HLA-I, while using proximity of PD-L1 to transferrin receptor as a control transmembrane protein (Fig.S2B). PD-L1’s proximity to HLA-I will enhance the likelihood of binding PD1 when HLA-I presenting antigen is engaged by the T cell receptor, thereby potentially contributing to PD-L1’s effectiveness as an immune checkpoint. Our data establish that the previously reported PD-L1 proximity to HLA-I proteins in lymphocytes^25^ is conserved in non-immune cells.

Gene ontology analysis of the 438 protein PD-L1 proteome revealed multiple biological pathways enriched among the PD-L1 proximal proteome, including an enrichment of proteins that function in the traffic of membrane proteins to and from the PM as well as among intracellular compartments (Fig.2G). These data agree with our functional data that PD-L1 traverses a number of intracellular compartments as it constitutively traffics between the PM and interior of cells.

### A role for PD-L1 in cell migration

Approximately 29% of the PD-L1 proximity map proteins are annotated to cytoskeleton organization, covering a broad range of both actin and microtubule biology (Fig.2H, Table S1). Because the actin and microtubule cytoskeletons have a role in the regulation of membrane protein traffic, this enrichment likely reflects, in part, the constitutive movement of PD-L1 among different cellular compartments. However, to explore the possibility that enrichment of cytoskeletal proteins in the PD-L1 proximity map also reflects a role for PD-L1 in cytoskeletal-dependent functions, we determined the effect of PD-L1 knockout (**KO**) on cell migration. PD-L1 KO was confirmed by western blotting (Fig.S3A). PD-L1 KO inhibited the migration of BEAS-2B cells in a scratched monolayer assay (Fig.3A, Fig.S3B). This finding is in line with previous reports that PD-L1 has a function in the regulation of cell migration^26^. Durvalumab, a therapeutic anti-PD-L1 antibody^27,28^, did not affect migration of the BEAS-2B cells, suggesting that the anti-tumor activity of antibody binding PD-L1 does not involve disruption of PD-L1’s control of migration (Fig.3B, Fig.S3B).

**Figure 3.**
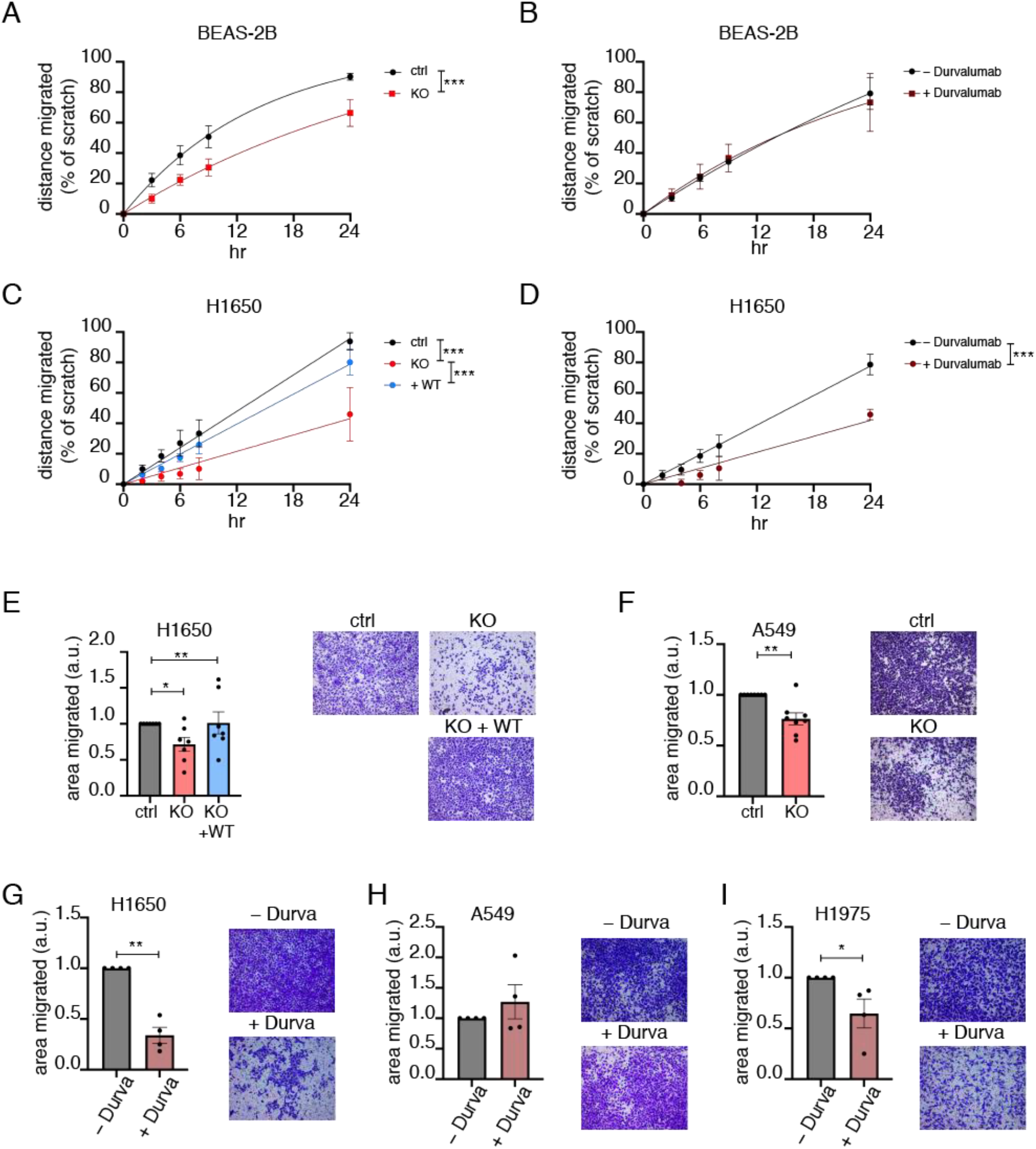
A role for PD-L1 in cell migration. (A-B) Quantification of scratch assay in (A) BEAS-2B PD-L1 KO cells, N=6, and (B) BEAS-2B treated with Durvalumab, N=4. For (A-B) rate constants are compared using extra sum-of-squares F test. Cells were in serum-free media for the duration of the assay. (C) Quantification of scratch assay in H1650 PD-L1 KO/rescue cells, N=4. Cells were in serum complete media for the duration of the assay. (D) Quantification of scratch assay in H1650 cells treated with Durvalumab, N=3. Cells were in serum-free media for the duration of the assay. For (C-D) difference in slope values is calculated using F-test. (E) Transwell migration in H1650 PD-L1 KO/rescue cells. One-way ANOVA and post-hoc Tuckey test. (F) Transwell migration in A549 PD-L1 KO cells. (G-I) Transwell migration in (G) H1650, (H) A549 and (I) H1975 cells treated with Durvalumab. For (F-I) paired student’s t-test was used on raw values. All error bars = SEM. * ≤ 0.05, ** ≤ 0.01, *** ≤0.001.

We next determined whether PD-L1 has a similar role in migration of cancer cells. PD-L1 KO and re-expression in H1650 cells were confirmed by western blotting (Fig.S3A). PD-L1 KO blunted migration of H1650 cells (an EGFR mutant cell line), an effect that was rescued by re-expression of PD-L1 (Fig.3C, Fig.S3C). Unexpectedly, Durvalumab inhibited migration of H1650 to the same degree as PD-L1 KO (Fig.3D, Fig.S3D). PD-L1 KO or anti-PD-L1 antibody did not affect H1650 cell proliferation, confirming the effects were on cell migration into the wounded monolayer (Fig.S3E-F). The different effects of anti-PD-L1 antibody on migration in non-transformed cells BEAS-2B and NSCLC cell line, H1650, suggest cell-context specific effects of αPD-L1 binding to PD-L1.

To determine whether the effect of anti-PD-L1 antibody on migration was specific to cancer cells, we studied another human NSCLC cell line, A549 cells. These cells do not migrate into a scratched monolayer; therefore, we monitored migration by determining penetration of cells into a filter disk. H1650 migration into the filter disk was inhibited by PD-L1 KO and rescued by re-expression of PD-L1, similar to the effect on H1650 migration measured by the scratch assay (Fig.3E). PD-L1 KO in A549 cells was confirmed by western blotting (Fig.S3A). PD-L1 KO in A549 cells reduced penetration into the filter, confirming a role for PD-L1 in migration in a second cancer cell line (Fig.3F).

Anti-PD-L1 antibody reduced H1650 cell migration into the filter disk, confirming the effect on migration of these cells into scratched monolayer (Fig.3G). However, as was the case for non-transformed BEAS-2B cells, anti-PD-L1 antibody had no effect on migration of A549 cells (Fig.3H). H1650 cells are EGFR mutant, whereas A549 are wild type for EGFR. Anti-PD-L1 antibody also reduced migration of another EGFR mutant NSCLC line, H1975, suggesting the effect of anti-PD-L1 antibody on migration is a feature of EGFR mutant cell lines, revealing the possibility of mutant EGFR cell context-dependent PD-L1 functions (Fig.3I).

### Anti-PD-L1 *antibody treatment inhibits turnover of mutant but not wild type EGFR*

A potential cell context-specific link between PD-L1 and EGFR, as suggested by the anti-PD-L1 antibody effect on migration of mutant EGFR cell lines was also supported by EGFR being a component of the PD-L1 proximal proteome (Fig.2F, Table S1). In addition to the enrichment of EGFR, ESCRT complex proteins, which have a role in regulation of intracellular trafficking of ligand activated EGFR, were also enriched in the PD-L1 proximal proteome (Fig 2F,H). We confirmed proximity to PD-L1 for EGFR and HGS (a subunit of ESCRT-0 complex) (Fig.S2B).

The ESCRT complex is involved in targeting membrane proteins to the lysosomes for turnover, and one of its best studied client proteins is ligand-activated EGFR^29,30^. Although proximity of PD-L1 to ESCRT proteins may reflect PD-L1 turnover, given the relative high enrichment of ESCRT0 and ESCRT1 proteins in the PD-L1 proximity map (Fig 2F) we explored the possibility of a link between PD-L1 and ESCRT-dependent function(s) beyond the targeting of PD-L1 for degradation.

Knockdown (**KD**) of PD-L1 in BEAS-2B cells (two-fold decrease in PD-L1 protein expression with KD) delayed ligand stimulated turnover of EGFR (Fig.4A, Fig.S4A). In both wildtype and PD-L1 KD cells, within 30 min of EGF stimulation, EGFR moved from a diffuse distribution on the PM to concentration in endosomes, demonstrating PD-L1 KD did not affect EGF-stimulated EGFR internalization. However, 60 and 90 min after 50ng/mL EGF stimulation, EGFR fluorescence was higher in the PD-L1 KD cells than control cells, demonstrating EGFR turnover was delayed by PD-L1 KD at a post-internalization step, perhaps in transport from the endosomes to lysosomes, a step requiring ESCRTs. PD-L1 KO, in a western blot assay for EGFR expression, also delayed EGF-induced receptor turnover, confirming a role for PD-L1 in controlling the post-stimulation half-life of EGFR (Fig.4B). αPD-L1 antibody did not affect the half-life of activated EGFR in BEAS-2B cells (Fig.4C). Thus, as was the case for migration of BEAS-2B cells, PD-L1 KO but not αPD-L1 antibody has effects on EGFR behavior in BEAS-2B cells.

**Figure 4.**
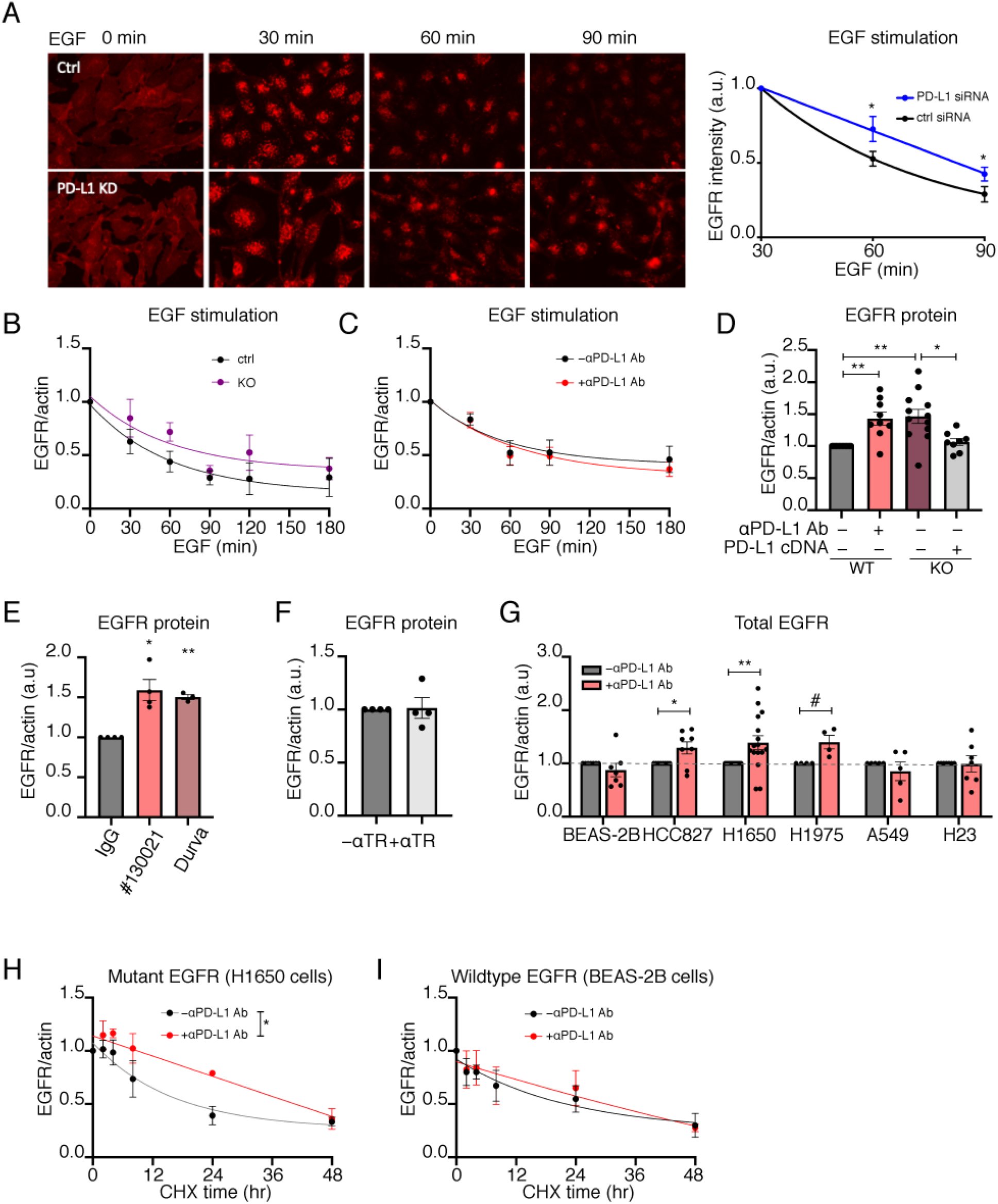
Anti-PD-L1 antibody treatment inhibits turnover of mutant but not wild type EGFR. (A) Representative images and quantification of EGFR in BEAS-2B cells with EGF stimulation, N=6. Paired student’s t-test at 60 and 90 minutes were calculated. (B-C) EGF-stimulated EGFR degradation in (B) BEAS-2B PD-L1 KO cells, N=4, and (C) BEAS-2B cells treated with αPD-L1 antibody, N=4. Rate constants are compared using extra sum-of-squares F test. (D) EGFR protein in WT, PD-L1 KO and PD-L1-reexpressing H1650 cells. One way ANOVA and post-hoc Tukey test. (E) EGFR protein expression in H1650 cells treated with reagent grade αPD-L1 antibody (clone # 130021, Novus Biologicals) and clinical Durvalumab as compared to isotype IgG control antibody. One-way ANOVA and post-hoc Tukey test. (F) EGFR protein in H1650 cells with anti-transferrin receptor antibody (aTfR) antibody treatment. One sample t-test. (G) EGFR protein expression in cell lines treated with αPD-L1 antibody. One sample t-test. (H-I) Quantification of EGFR degradation in (H) H1650, N=3, and (I) BEAS-2B, N=3, in the presence of cycloheximide. Rate constants are compared using extra sum-of-squares F test. For (B-I) Western blot quantifications of EGFR protein normalized to actin expression are shown. All error bars = SEM. * ≤ 0.05, ** ≤ 0.01, *** ≤0.001, # = 0.051.

In H1650 cells, both PD-L1 KO and αPD-L1 antibody increased mutant EGFR protein amounts by about 50%, and re-expression of PD-L1 fully rescued the effect of the knockout (Fig.4D, Fig.S4B). Durvalumab and lab reagent anti-PD-L1 antibody (clone# 130021, Novus Biologicals), which bind different extracellular epitopes of PD-L1, were equally effective at increasing mutant EGFR protein amounts, whereas an antibody targeting the extracellular domain of the transferrin receptor, another PM protein, did not affect the amount of EGFR protein (Fig.4E-F, FigS4C,D). Analyses of a panel of NSCLC lines demonstrated that the effect of αPD-L1 antibody on turnover of the ligand-stimulated EGFR protein amount was specific to cells harboring mutant EGFR (Fig.4G). αPD-L1 antibody caused a decrease turnover, evidenced by increase in the amount of EGFR in three different EGFR mutant lines, whereas the amounts of EGFR in two KRAS mutant lines and a non-transformed line were unaffected. The effect is not linked to a specific EGFR mutation because the EGFR mutant lines harbor different activating mutations, nor was the effect of αPD-L1 antibody associated with differences in PD-L1 expression among the cell lines (Fig.1D). Furthermore, anti-PD-L1 antibody treatment did not affect the distribution nor total amount of either PD-L1 or the transferrin receptor in EGFR mutant cells (Fig.S4E, F).

To directly monitor the effect of anti-PD-L1 antibody on EGFR half-life, we compared EGFR turnover in cells treated with cycloheximide to inhibit translation. In these conditions the time dependent reduction of mutant EGFR reflects its turnover. anti-PD-L1 antibody significantly increased mutant EGFR half-life without an effect on wildtype EGFR (Fig.4H-I, Fig.S4G). These data also demonstrate the increased mutant EGFR protein amount is not due to an increase in translation. In addition, anti-PD-L1 antibody did not alter mutant EGFR mRNA amounts, supporting the conclusion that anti-PD-L1 antibody modulates mutant EGFR protein levels by increasing its half-life (Fig.S4H, I). The extension of mutant EGFR half-life by anti-PD-L1 treatment did not augment EGFR signal transduction measured by changes in gene transcription (Fig.S4J).These data on EGFR turnover provide additional support for the hypothesis that EGFR mutant cells have a heightened sensitivity to perturbation of PD-L1 by anti-PD-L1 antibody binding, while PD-L1 KO, a more severe disruption of PD-L1, impacts both WT and mutant EGFR cell lines.

### Anti-PD-L1 antibody treatment affects extracellular vesicles

ESCRTs, in addition to having a key role in targeting EGFR (and other proteins) for lysosomal degradation, are also involved in the formation and secretion of extracellular vesicles (EVs)^31^. To assay EV formation, we used mass spectrometry to identify proteins in EVs isolated from the growth media of BEAS-2B, A549 and H1650 cells treated with or without anti-PD-L1 antibody for 48hr. We also analyzed the whole cell extract collected from the same cells. To limit the analysis to most highly enriched EV proteins, we focused on proteins with EV to whole cell extract (WCE) ratio in the top quartile for each respective cell line. Although there was some variation, the lists largely overlapped among the three cell lines (Fig.5A, Table S2). The biological pathways enriched among the EV proteins of the 3 cell lines were similar and the biological pathways of the identified proteins were congruent with reported functions of EVs, including exosome formation, cell adhesion and migration as well as immune cell extravasation^32,33^ (analysis of H1650 EV is shown Fig.5B).

**Figure 5.**
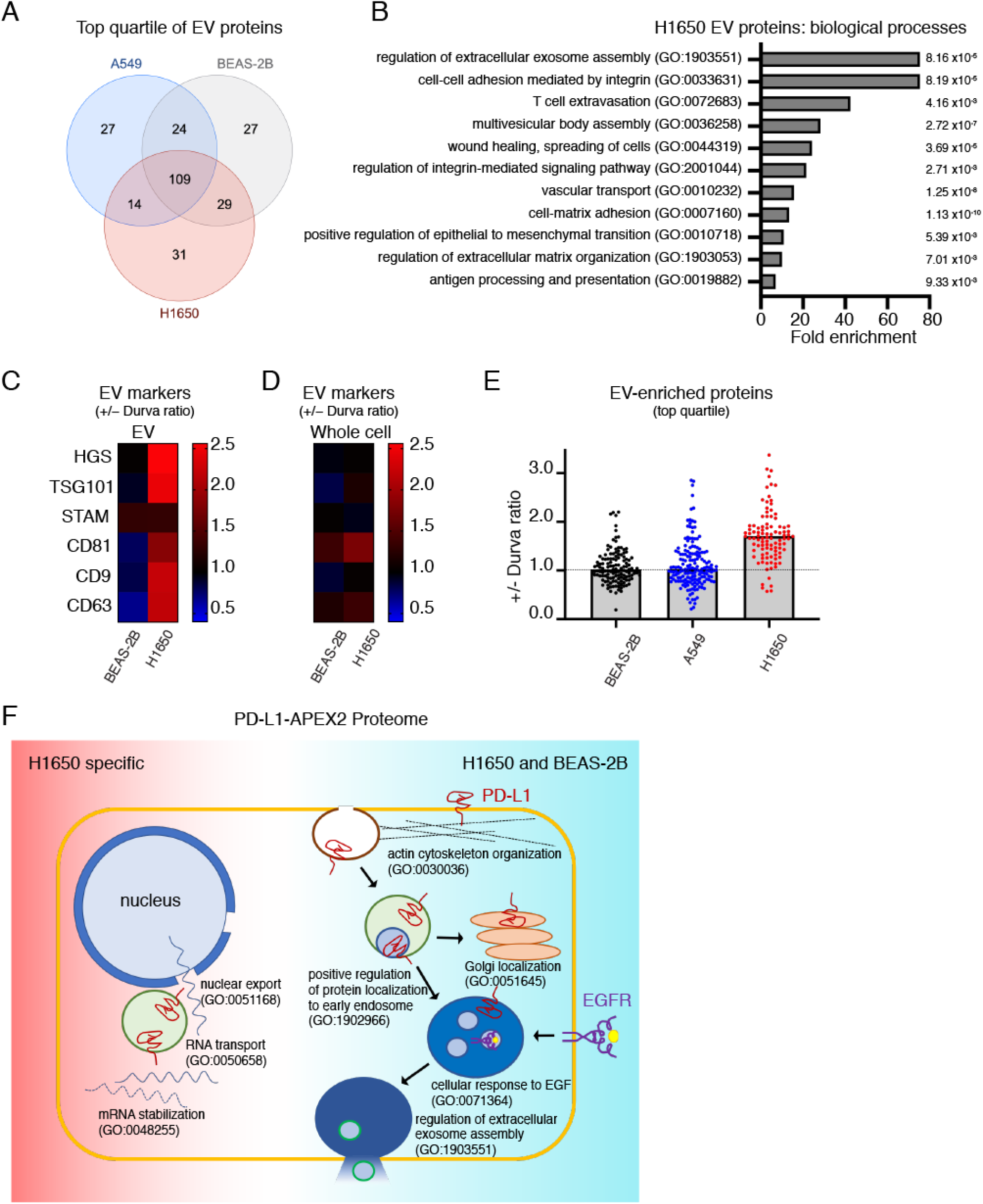
Anti-PD-L1 antibody treatment affects extracellular vesicles. (A) Venn diagram of EV proteins with the highest intensities (top quartile) in each of the cell lines. (B) Biological processes representing proteins that are most enriched in the EVs of H1650 cells. Fold enrichment and FDR for the pathways are shown. (C) Heatmap of Durvalumab-induced fold change of extracellular vesicle marker proteins expression in (C) EV fraction and (D) whole cell lysate in BEAS-2B and H1650. (E) Durvalumab-induced fold change of extracellular vesicle-enriched proteins (top quartile for each cell line) in BEAS-2B, A549 and H1650. For (C-E) cells were treated with Durvalumab. Average +/− Durvalumab value for N=2 is shown. (F) Schematic of H1650 PD-L1-APEX2 proximity map shared with BEAS-2B and unique to H1650.

Next, we analyzed if anti-PD-L1 antibody affects EV secretion or content. As a general measure of EV formation, we first examined a set of 6 canonical EV proteins: HGS, TSG101, STAM, CD81, CD9 and CD63^34^ (Fig.5C). Durvalumab did not affect the amounts of these proteins in EVs isolated from BEAS-2B cells, whereas Durvalumab increased these proteins, except for STAM, in EVs isolated from H1650 (Fig.5C). anti-PD-L1 antibody did not affect the amounts of these proteins in whole cell extracts (Fig.5D).

Consistent with the effects of anti-PD-L1 antibody on the 6 canonical EV proteins, anti-PD-L1 antibody treatment caused a general increase of proteins detected in EVs isolated from H1650 conditioned media and elicited no change in BEAS-2B or A549 (Fig 5E). These data support αPD-L1 antibody increasing EV biogenesis and or protein packaging selectively in EGFR mutant cells.

### Anti-PD-L1 Proximity Proteome in EGFR mutant cells

Our functional data support the hypothesis that EGFR mutant cells are uniquely sensitive to anti-PD-L1 treatment. To determine whether this difference is revealed by a difference in PD-L1 protein in EGFR mutant cells, we repeated PD-L1-APEX2 proximity labeling in H1650 cells. There was considerable overlap in the PD-L1 proximity proteome between WT and EGFR mutant cells revealed at the level of biological pathways, including significant enrichments of proteins annotated to the control of membrane protein traffic pathways and to various aspects of cytoskeleton biology (Fig.5F). At the individual protein level, there is greater than 60 percent correspondence in the PD-L1 proximal proteome between the WT and mutant EGFR cell lines, including proximity of PD-L1 to EGFR and ESCRT components (Table S3). These data suggest that a difference in PD-L1 localization between wild-type and mutant EGFR does not account for the sensitivity of EGFR mutant cells to αPD-L1 antibody treatment. Interestingly, PD-L1 proximity proteome in H1650 uniquely revealed proteins involved in nuclear export as well as RNA stability, which could be related to its role in regulation of RNA stability^35^.

## Discussion

PD-L1 is a transmembrane ligand for the PD1 immune checkpoint receptor^21,22,36,37^. Tethering PD-L1 to the membrane provides spatial constraint on its immune checkpoint activity because the PD1-bearing cell must be close enough for PD-L1 on the surface of the expressing cell to bridge the distance between cells. Our proximity mapping reveals a second feature of spatial control of PD-L1 checkpoint activity. We identified HLA-I proteins are in close proximity to PD-L1 in non-transformed human lung epithelia cells and mutant EGFR NSCLC cell line (Tables S1 & S3). This juxtaposition of HLA-I and PD-L1, formed in the absence of HLA-I binding to the TCR, ensures that PD-L1 is poised to engage PD1 once HLA-I has engaged the TCR, perhaps enhancing the efficiency of PD-L1 to attenuate TCR activation. This PD-L1 to HLA-I proximity is similar to PD1 to TCR proximity on the T cell^38–40^. Our findings extend the previous observation of PD-L1 adjacent to HLA-I in leukocytes by demonstrating this spatial control is not solely restricted to immune-to-immune cell interaction and that it exists in the absence of TCR-HLA-I engagement.

Because PD-L1 is a transmembrane protein, its PM density can be modulated by controlling membrane protein traffic. Our kinetic analyses demonstrate that the amount of PD-L1 is actively maintained by constitutive cycling between the interior of cells and the PM (Fig.1). Proteins that regulate membrane protein traffic from and to the PM were enriched in the PD-L1 proteome, supporting our results that PD-L1 is constitutively transiting multiple membrane compartments rather than being statically localized to the PM and or a single intracellular site (Fig.2,Table S1). Others have previously probed the regulation of PD-L1 trafficking. Most notably, it has been shown that CMTM6 plays a key role in controlling the fate of internalized PD-L1^11,12^. CMTM6 KO results in increased degradation of PD-L1, reducing both PM PD-L1 and total cellular PD-L1. Our results established growth factor regulation of constitutive PD-L1 recycling as an additional mechanism for regulation of PD-L1 PM density (Fig.1). Growth factor regulation, mediated by AKT activity, is independent of CMTM6 and IFNγ regulation of total cellular PD-L1. Both CMTM6 and AKT-dependent control are acute modulators of PD-L1 density, acting more rapidly than IFNγ transcriptional regulation, which could be advantageous in defending against normal tissue damage at a site of an active immune response, likely non-pathologic activity of PD-L1. AKT control of PD-L1 traffic was similar in non-transformed and transformed cells and it was unaffected by whether the driver oncogene was mutant EGFR or KRAS, indicating this is a fundamental feature of PD-L1 biology and not specific to cancer cells or specific driver mutation.

Regulation of PD-L1 recycling has an advantage compared to CMTM6 in the control of PD-L1 PM density because the changes are rapidly reversible since the location of PD-L1 not the total amount are modulated. Reduced growth factor signaling slows recycling of PD-L1, shifting PM PD-L1 to intracellular sites while maintaining constitutive cycling, albeit at a reduced rate (Fig. S1C). Restoration of growth factor AKT activation redistributes intracellular PD-L1 to the PM by increasing recycling without requiring new synthesis of PD-L1, whereas reversal of CMTM6 depletion of PD-L1 requires new synthesis to overcome the effect of increased PD-L1 degradation.

Many studies have highlighted PD-L1 functions that are in addition to its role as an immune checkpoint ligand, including regulation of cell growth, EMT, and resistance to cellular stresses such as chemotherapy and IFN-mediated cytotoxicity^7,8,41^. Biological pathways enriched among the PD-L1 proximal proteome support the conclusions of many of those previous studies. Our studies of PD-L1 within different cell contexts (non-transformed, EGFR mutant and KRAS mutant cells) advances the prior studies by revealing novel cell-context dependencies for PD-L1 role in cell migration as well as the formation and or release of exosomes.

PD-L1 has been shown to have a role in migration of cell lines with non-EGFR driver oncogenes^26^. Those studies relied on genetic manipulation of PD-L1. We confirm and extend those studies by showing genetic ablation of PD-L1 inhibits cell migration in non-transformed, EGFR mutant and KRAS mutant cell lines (Fig.3). Strikingly, we found that anti-PD-L1 antibody treatment inhibited migration of EGFR mutant cells but had no effect on KRAS mutant (wild type EGFR) or non-transformed cells. We do not know why EGFR mutant cells are sensitive to anti-PD-L1 antibody in a way that other cells are not. In addition to possible changes in cell physiology arising from the constitutive EGFR signaling of mutant EGFR, wildtype and mutant EGFR have been described to have largely dissimilar downstream signaling^42,43^. Our data suggest that the cellular context created by oncogenic EGFR signaling may make EGFR mutant cells more susceptible to anti-PD-L1 antibody-mediated signaling changes. In support of that view, a recent study showed PD-L1 induces phospholipase C-γ1 (PLC-γ1) activation only in the presence of activated EGFR^44^, although additional experiments are required to determine whether PLC-γ1 has a role in anti-PD-L1-mediated effects in mutant EGFR cells.

ESCRT proteins control the turnover of transmembrane proteins and extracellular vesicle formation (EV) by inducing the intralumenal budding of vesicles from the limiting membrane of endosomes to form multivesicular bodies (MVBs)^29,45,46^. The ESCRT pathway is key in the negative regulation of EGFR signaling by controlling EGFR degradation^47^. In agreement with previous studies identifying PD-L1 proximity with EGFR^48^, and ESCRT accessory protein ALIX regulating PD-L1 sorting into intralumenal vesicles^15^, we identified EGFR as well as several ESCRT complex proteins as components of the PD-L1 proximal proteome (Table S1). Steady-state PD-L1 proximity to the ESCRT machinery might reflect normal turnover of PD-L1 by lysosomal degradation or EV secretion^11,13,15,49^. However, our analyses strongly support the hypothesis that PD-L1 is functionally associated with the ESCRT pathway beyond being a client for MVB sorting. Anti-PD-L1 treatment had two effects on ESCRT-dependent functions in EGFR mutant cells: turnover of constitutively active EGFR was significantly reduced and biogenesis/secretion of EVs was significantly increased.

Notably, the effects of anti-PD-L1 treatment on ESCRT pathway functions, like the effect on cell migration, were only observed in EGFR mutant cells. Genetic ablation of PD-L1 in wild type EGFR cells mimics the effects of anti-PD-L1 treatment on EGFR mutant cells, suggesting a more significant perturbation is required to disrupt cell-intrinsic PD-L1-dependent functions in wild type EGFR cells. We do not understand what it is about constitutive EGFR signaling that sensitizes cells to the effects of anti-PD-L1 therapy.

The effect of antibody binding to PD-L1 has not been extensively studied in the context of PD-L1 reverse signaling. Our data suggest antibody binding in mutant EGFR cells results in a loss of function of PD-L1 as it phenocopies the effect of PD-L1 KO on EGFR expression, cell migration and EV secretion. Future studies are required to determine whether the effects of anti-PD-L1 antibody binding are due to a perturbation in PD-L1 transmembrane signaling or disruption in PD-L1 binding with partner proteins, which are responsible for signaling changes. Recent studies have identified PD-L1 signaling motifs and binding partners that result in broad transcriptional and signaling changes within the cells^8,44,50^.

The effects of anti-PD-L1 antibody on EGFR mutant cells we discovered might be informative for understanding the increased resistance of EGFR mutant tumors to anti-PD-L1 therapy^5,6,51^. In particular, the anti-PD-L1 antibody-induced increase in EVs from mutant EGFR cells could have broad ranging effects on the tumor microenvironment and tumor immune landscape that could contribute to resistance to immunotherapy^52,53^. PD-L1 was not detected by mass spectrometry in EVs from any of the cell lines, therefore our data do not speak to the possibility that PD-L1 on EVs serves to perturb immune activity or that it affects anti-PD-L1 therapy as reported^52^.

PD-L1 expression on immune cells has an important immune-modulatory role^50,54–56^ that will also be affected by anti-PD-L1 therapy. For example, PD-L1 on dendritic cells (DCs) is necessary for efficient homing of dermal DCs to the lymph nodes through modulation of chemokine receptor signaling and actin remodeling^50^. Our finding that the effects of anti-PD-L1 are cell-context dependent raise the possibility that anti-PD-L1 effects in immune cells extend beyond blocking PD1 engagement.

### Limitations

As a hypothesis generating study our work provides a foundation for future work, in particular studies designed to better understand the specific sensitivity of mutant EGFR cells to anti-PD-L1 antibody. A limitation of our work is that all studies were performed in cultured cells. However, we are technically constrained in this regard because there are no syngeneic mouse mutant EGFR cell lines that can be used for *in vivo* testing of hypotheses emerging from our *in vitro* data in an immune competent mouse.

## Materials and Methods

### Cell lines and culture

BEAS-2B (non-tumorigenic), A549 (KRAS mutant), and H23 (KRAS mutant) were purchased from ATCC. H1650, HCC827, HCC4006 and PC9 (all EGFR Ex19Del mutant) were a gift from Dr. Rafaella Sordella (Cold Spring Harbor Laboratory). H1975 (EGFR L858R/T790M double mutant) were a gift from Dr. Brandon Stiles (Weill Cornell Medicine). All primary human samples were obtained and used in accordance with an approved IRB protocol. Primary tumor cells were processed to obtain single cell suspensions as described previously^57^ and plated onto glass-bottom dishes and analyzed 3-5 days later. All cells were cultured in DMEM media supplemented with 10% FBS and 1% penicillin-streptomycin at 37°C and 5% CO_2_. Where indicated, cells were treated with serum-free DMEM media supplemented with 0.02M HEPES and 0.03M NAHCO_3_.

### Cell treatments

For PD-L1 PM to total measurements, cells were plated on glass bottom dishes in full growth media and assayed the next day. Cells were switched to either serum-free media, serum free media supplemented with growth factors or fresh full growth media for 4 hours. EGF was used at 8nM, IGF at 10nM and insulin at 10nM. For Akt inhibition experiments, cells were incubated in full growth media with 5 μM MK2206 or DMSO for 4 hours. Interferon gamma treatment was performed at 40 ng/mL for 24 hrs, and CMTM6 shRNA knock-down was induced by doxycycline at 1 μg/mL for 48hrs. For antibody treatment experiments, cells were treated with specified antibody for 7 hr at a final concentration of 10 μg/mL in serum free DMEM media. For protein turn-over experiments cycloheximide was used at 10 μg/mL. For stimulated EGFR degradation, cells were serum starved for 4 hr and treated with EGF at 50ng/mL for indicated timepoints respectively, while antibody was used at 10 μg/mL and maintained for the duration of the experiment. For EGFR signaling experiments, cells were incubated with antibody in serum free media for 7 hr and stimulated with 0.5 ng/mL EGF for specified timepoints. For migration assays, cells were pre-incubated with Durvalumab antibody for 5 days before being plated for experiment.

### Extracellular vesicle and whole cell extract preparation for mass spectrometry

Cells were adjusted to media supplemented with 10% exosome free FBS for 1 passage and plated at 60% confluency in exosome free media. After 48 hours of Durvalumab treatment (10 μg/mL), cell media was collected for extracellular vesicle isolation and cells were lysed in 1x RIPA buffer with protease/phosphatase inhibitors. Extracellular vesicles were isolated by sequential ultracentrifugation (12000g for 10 min to discard cell debris, 20000 g for 20 min to discard microvesicles, 100000 g for 70 min twice to isolate and wash the EVs) and resuspended in a final volume of 100 μL of RIPA buffer with protease/phosphatase inhibitors. Cell extract and EV lysates were quantified for protein concentration. A final amount of 100 μg of WCE, and equi-volume of EVs approximating 10 μg of protein in each sample was submitted for mass spectrometry analysis.

### PD-L1-APEX2 labeling

APEX2 proximity assay was performed as described previously^58^. Briefly, BEAS-2B cells stably expressing PD-L1 APEX2 were grown until confluency. Cells were preincubated with biotin phenol in full growth media for 30 minutes and hydrogen peroxide was added for 1 minute (except in negative control). H1650 PD-L1-APEX2 stably expressing cells were treated similarly with the exception of biotin phenol concentration (500 μM for BEAS-2B and 2.5 mM for H1650). Cells were washed three times with ice cold quenching solution (10 mM sodium ascorbate, 5 mM Trolox and 10 mM sodium azide solution in PBS) and cells were lysed in 1x RIPA lysis buffer with protease/phosphatase inhibitors. Protein concentration was measured, and biotinylated proteins were pulled down using streptavidin-agarose beads overnight at 4°C. Beads were washed (twice with RIPA lysis buffer, once with 1 M KCl, once with 0.1 M Na2CO3, once with 2 M urea in 10 mM Tris-HCl,pH 8.0, and twice with RIPA lysis buffer) and pull-down proteins were submitted for mass spectrometry.

### Mass spectrometry data analysis

BEAS-2B PD-L1-APEX2 data were analyzed in two different ways. In one, proteins, in which protein intensity in the experimental sample was two-fold or greater than in the control sample, were identified to be in the proximity of PD-L1. Data from 5 independent experiments were analyzed individually and the final PD-L1 proximity map was determined as those proteins satisfying the criteria for PD-L1 proximity in each of the five experiments, yielding 438 PD-L1 proximity proteins. This set of proteins was used for further analyses, including biological processes pathway analysis via geneontology.org (GO database released 11-16-2021)^59^. The data were also analyzed by performing a t-test on Log10 transformed spectral intensity values between labeled and negative control samples (-H_2_O_2_) on the set of 1647 proteins common to all five experiments. Significant differences between the experimental and control were determined using Benjamini-Hochberg correction for multiple testing. Fold change was calculated as ratio of average spectral intensity in labeled samples over average in negative control samples (as shown in Fig 2F). The two different methods yielded essentially the same group of proteins. For H1650 PD-L1-APEX2 data, proteins with intensities 2-fold or higher in experimental sample over control in all 3 replicate experiments were identified as PD-L1 proximal.

For EV mass spectrometry analysis, for each protein its intensity in EV was divided by the corresponding intensity in whole cell extract (WCE). Proteins that were not detected in either sample were excluded. The remaining proteins were rank ordered based on the EV/WCE ratio. This analysis was done for both conditions (+/− Durvalumab) and both biological replicate experiments (4 samples total for each cell line). Proteins falling in the top quartile in 3 out of 4 samples in each cell line were used for downstream analyses.

### Migration and proliferation

For scratch assays 35K cells were plated on each side of the Ibidi 2-well scratch assay chambers. Cells were allowed to attach for 24 hrs and wound healing was assayed in complete or serum-free media for KO/rescue cells and Durvalumab treated cells respectively at specified timepoints. Cells were imaged in bright-field on Leica Microscope. 9 images of each sample were taken at each timepoint, and three distances were measured in each image using Image J. For migration assay, 30k cells were plated on the top chamber of transwell-migration plates with 8 micron pore size in serum free media. Bottom chamber was filled with DMEM with 10% FBS and cells were allowed to migrate for 24 hrs. Cells were then fixed in 3.7% formaldehyde for 15 minutes, washed and then stained in Crystal Violet for 20 minutes. Unmigrated cells were scraped off with damp cotton swab and membrane was cut out and mounted on slides. Migrated cells were imaged on upright microscope (Zeiss Axioplan) with 10x objective and analyzed via ImageJ. For proliferation assay, 5k cells were plated in triplicate 96 wells (day 0) and MTT was used to quantify viable cells on days 1-5. Paired t test was performed on raw transwell migration data, and slopes of nonlinear regression lines were used to calculate significance for scratch and MTT assays.

## Acknowledgements

We are grateful to Dr. Raffaella Sordella (Cold Spring Harbor Laboratories) for generously sharing EGFR mutant cell lines with us. We thank Astra Zeneca for the gift of Durvalumab antibody. We thank Dr. Alberto Benito and Dr. Kristy Brown for help with EV purification. We thank Rosemary Leahey for assistance with imaging studies. We thank the Optical Microscopy & Image Analysis Core at Weill Cornell Medicine for assistance with confocal microscopy. We thank Guoan Zhang and the Weill Cornell Medicine Proteomics and Metabolomics Core for assistance with whole cell and EV proteomics. We thank Dr. Jeremy Dittman, Dr. Geoffrey Markowitz, Dr. Lucie Yammine, Dr. Anuttoma Ray, Jennifer Wen, and Emma Johnson for helpful discussions and critical reading of the manuscript.

The project was supported in part by NCI UG3 CA244697 (NKA, TEM), the Yoram Cohen family foundation (NKA), Vicky and Jay Furhman family fund (NKA) and a WCM Meyer Cancer Center pilot grant (NKA,TEM). TEM is also supported by DOD LC180227. EY is supported by a diversity supplement to CA244697.

## Author contributions

AL designed and performed the experiments, analyzed the data, and wrote the manuscript. EY designed experiments, generated the KO cells, and edited the manuscript.

PC and ND performed and assisted with APEX2 mass spectrometry experiments and analysis NKA conceptualized the project, contributed to experimental design and interpretations, and edited the manuscript.

TEM conceptualized the project, contributed to experimental design, performed analysis and interpretation, and wrote the manuscript.

## Notes

### Competing Interest Statement

The authors have declared no competing interest.

### Summary of Updates

The text has been revised to clarify data presentation and some additional experiments have been added to bolster the conclusions, although the overall conclusions of the revised manuscript are in line with the original version.

