## Supplemental Figures for "Proximity proteome mapping reveals PD-L1-dependent pathways disrupted by anti-PD-L1 antibody specifically in EGFR-mutant lung cancer cells"

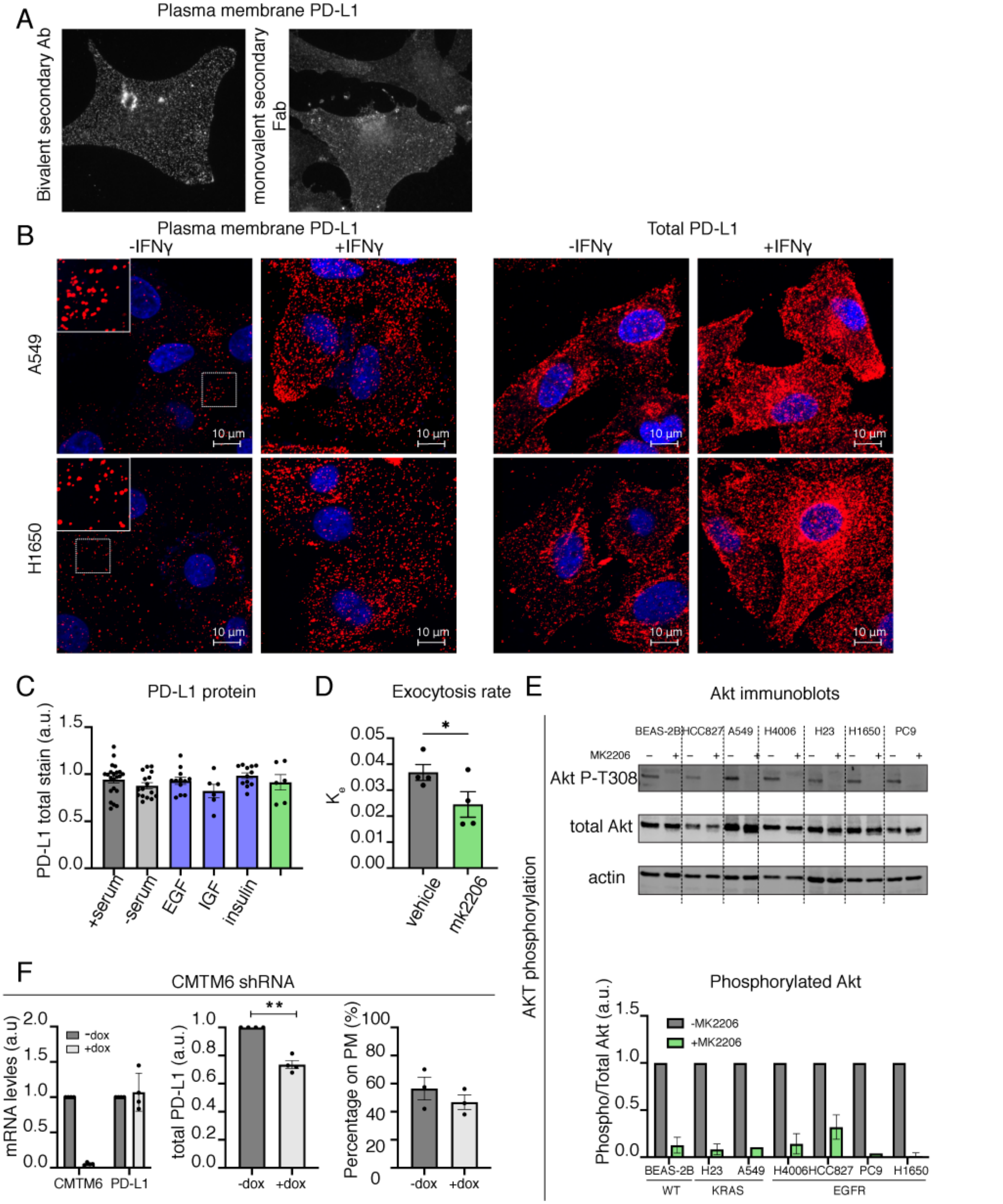


**Supplementary Figure 1. PD-L1 is dynamically maintained on the plasma membrane of cells.** (A) Epifluorescent images of BEAS-2B cells treated with IFNγ and stained for PM PD-L1 using bivalent secondary antibody or monovalent secondary Fab. (B) Representative confocal images of human lung cell lines with or without IFNγ treatment and immunostained for PM and total PD-L1. Red: PD-L1. Blue: Hoechst. Sum intensity of z-stacks is projected. Fluorescence intensity is equally scaled across panels for each cell line. Insets in -IFNγ PM are enhanced for better visualization. (C) Total PD-L1 protein expression in BEAS-2B cells with growth factor treatment or Akt inhibition. One-way ANOVA. (D)

Exocytosis rate constants in BEAS-2B cells incubated with vehicle (DMSO) or Akt inhibitor. Paired student's t-test. (E) Immunoblots showing Akt T308 phosphorylation in cells treated with vehicle (DMSO) or Akt inhibitior. Corresponding quantification of phospho-T308/total Akt is shown in the graph. (F) CMTM6 and PD-L1 mRNA expression, PD-L1 protein expression and distribution in BEAS-2B expressing doxycycline inducible CMTM6 shRNA. One sample and unpaired student's t-tests. All error bars = SEM. * ≤ 0.05, ** ≤ 0.01, *** ≤0.001

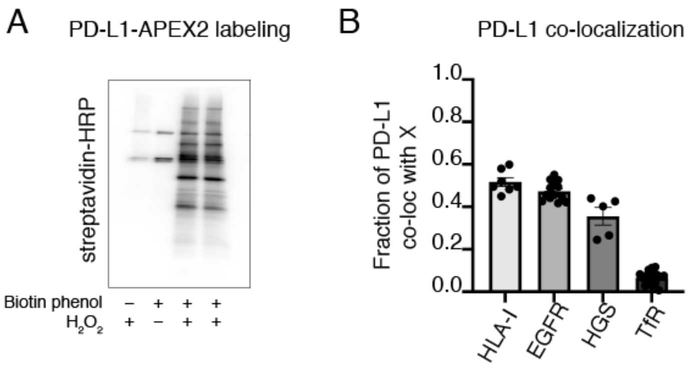


**Supplementary Figure 2. PD-L1 proximity mapping**
(A) Representative immunoblot of biotinylated proteins in labeled and control BEAS-2B cells. (B) PD-L1 co-localization with specified proteins quantified by fraction of PD-L1 intensity co-localized with each target protein. Each point is a collected from single plane images of individual cells under confocal microscopy.

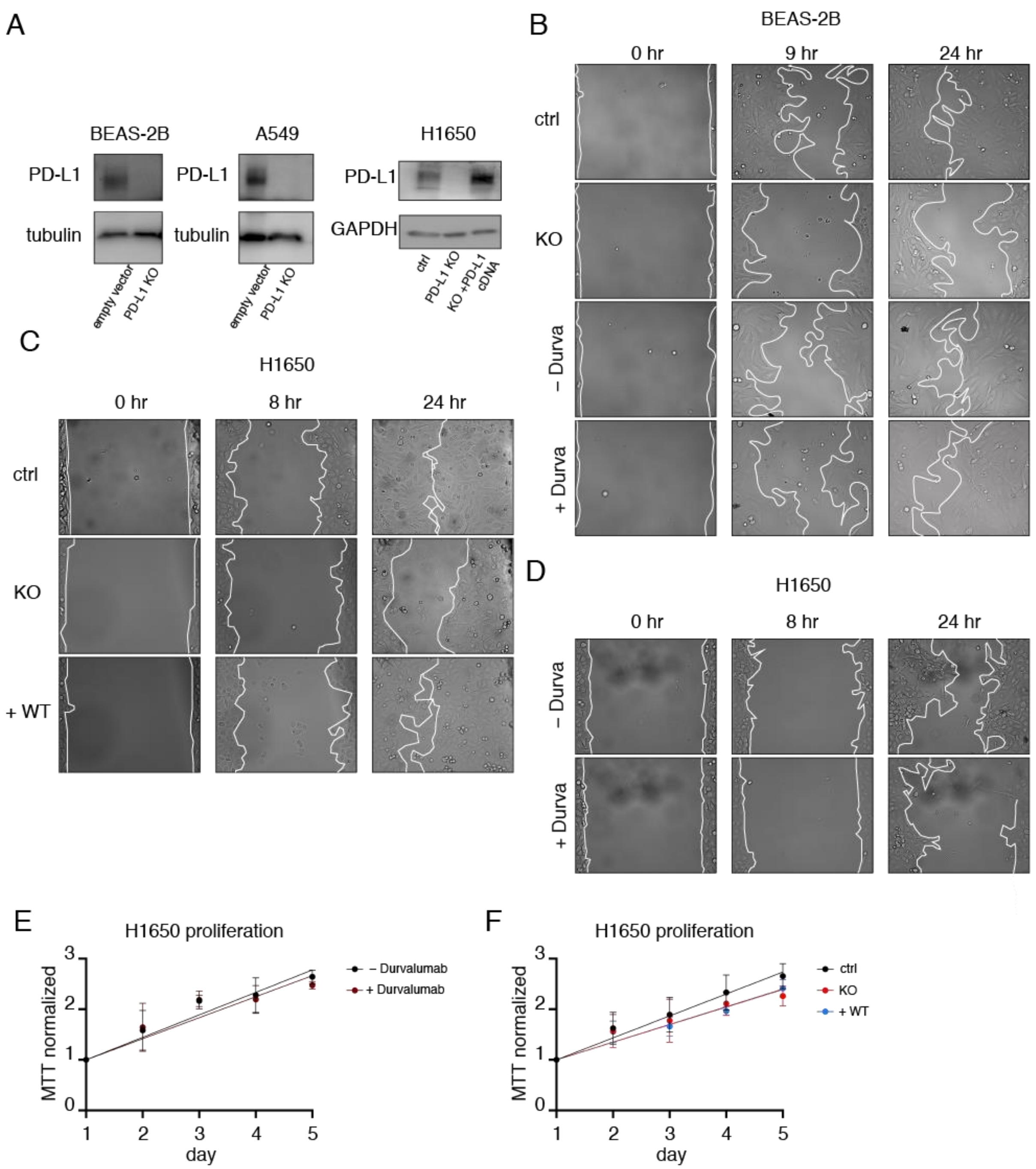


**Supplemental Figure 3. A role for PD-L1 in cell migration**
(A) Immunoblots of PD-L1 in control and PD-L1 KO cell lines. Tubulin is used as loading control. (B) Timelapse images of scratch assay at 0, 8 and 24 hr in BEAS-2B cells. (C-D) Timelapse images of scratch assay at 0, 8 and 24 hr in (C) H1650 PD-L1 KO rescue cells and (D) H1650 cells treated with Durvalumab. For (B-D) Leading edge of the cells are marked in white. Cells were in (B, D) serum-free media and (C) serum-complete media for the duration of the assay. (E-F) Cell proliferation as measured by MTT assay in (E) H1650 cells treated with Durvalumab, N=2, and (F) H1650 PD-L1 KO rescue cells, N=3. For (E-F) difference in slope values is calculated using F-test.
All error bars = SEM. * ≤ 0.05, ** ≤ 0.01, *** ≤0.001

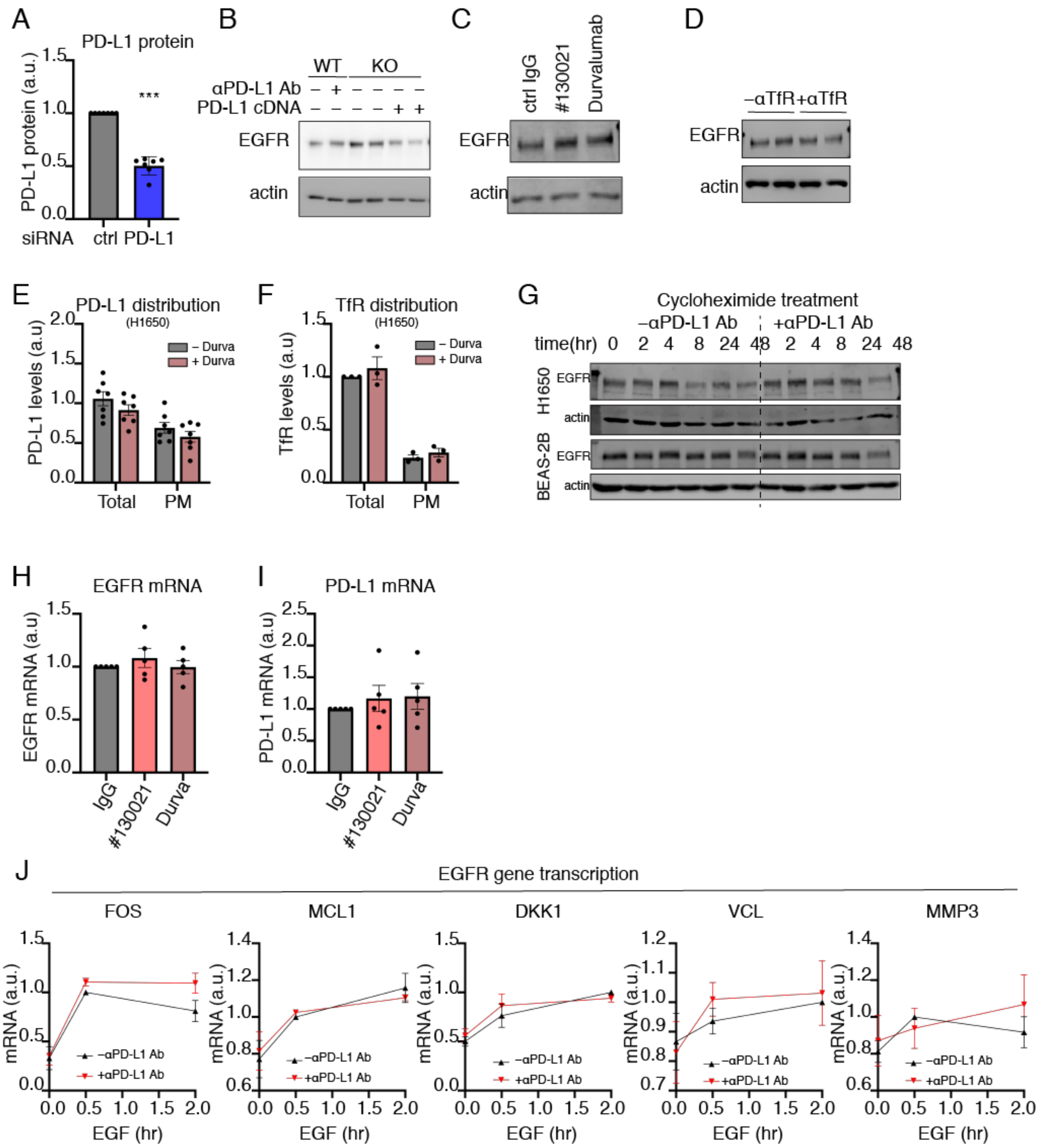


**Supplemental Figure 4. Anti-PD-L1 antibody treatment inhibits turnover of mutant but not wild type EGFR**

(A) PD-L1 protein expression in BEAS-2B cells transfected with control and PD-L1 siRNA. One sample t-test. (B-D) Immunoblots of EGFR expression in (B) H1650 PD-L1 KO/rescue cells and H1650 cells treated with (C) PD-L1 antibody and (D) TfR antibody. (E-F) Distribution of (E) PD-L1 and (F) TfR in H1650 cells treated with Durvalumab. Unpaired multiple t-tests. G) Immunoblots of EGFR degradation in BEAS-2B and H1650 treated with cycloheximide in serum complete media. Actin is used as loading control. (H-I) mRNA expression of (H) EGFR and (I) PD-L1 in H1650 cells treated with αPD-L1 antibodies. (J) EGF stimulated gene transcription in H1650 cells treated with αPD-L1 antibody. qRT-PCR mRNA quantification of EGFR target genes is shown, N=3. All antibody treatments are. All error bars = SEM. * ≤ 0.05, ** ≤ 0.01, *** ≤0.001
