## Supplemental Figures for "Proximity proteome mapping reveals PD-L1-dependent pathways disrupted by anti-PD-L1 antibody specifically in EGFR-mutant lung cancer cells"

Stable cell lines:

Stable cell lines were generated via lentiviral infection using Lenti-X packaging system. Cells were either antibiotic selected via blasticidin (5 μg/mL) or puromycin (3 μg/mL) or sorted for expression of protein of interest via FACS. PD-L1 KO cells were generated via CRISPR/CAS9 with both targeting sequences (guide1: 5’-CGGGAAGCTGCGCAGAACTG-3’; guide2: 5’ -TCTTTATATTCATGACCTAC-3’). Live cells were stained with PD-L1 antibody and KO cells were sorted by FACS. Bulk population of KO cells was verified for PD-L1 KO via Western Blot. CMTM6 stable inducible shRNA knockdown cells were generated with one shRNA targeting sequence (5’-TTCAAAGAAGTAGAGGCCTCCA-3’) and mRNA knock-down was confirmed via qRT-PCR.

Microscopy:

Cells were plated on glass-bottom dishes prior to all experiments. After corresponding experimental treatments, cells were washed and fixed in 3.7% formaldehyde for 5 minutes. Cells were stained with primary and fluorescent secondary antibodies for proteins of interest on the plasma membrane without permeabilization. Saponin (0.25 mg/mL) was used to permeabilize the cells for total protein stain. Cell nuclei were labeled with Hoechst stain. For quantifications, images were acquired on an inverted microscope with a 20× air objective or a 40x oil objective (Leica Biosystems DMI6000-B). Images were analyzed on MetaMorph image processing software and the background corrected average intensity under each cell’s area was used to quantify protein amount. The values were normalized to the average of replicate coverslip dishes in control condition in each experiment to allow for comparison across experiments. Unpaired two tailed Student’s t test or paired ratio t-test were performed on unnormalized raw data, and one sample t test was used on normalized data respectively as indicated. For comparison of three and more groups, one-way ANOVA was performed on raw data with post-hoc Tuckey test where indicated. For PD-L1 stain images, Z stack of the cells was acquired using Zeiss Confocal microscope (LSM880) with Airyscan on a 40x oil objective.

Exocytosis assay:

Live cells were incubated with a saturating concentration of PD-L1 antibody (50 μg/mL) for indicated times, washed extensively with PBS, and then fixed. All cells were permeabilized and stained with a secondary antibody to visualize all antibody bound to the cells. Separate set of dishes was used to quantify total PD-L1 stain in fixed cells in each experiment. Cells were imaged and quantified as described above. All values were normalized to the plateau, which is defined as the average of values at 60 and 120min in each experiment.

Colocalization analysis:

For co-localization analysis cells were fixed and stained for total PD-L1 and the protein of interest with primary antibody followed by Cy3 and Alexa488-labeled secondary antibodies. Cells were imaged on Zeiss Confocal microscope (LSM880) with Airyscan on a 40x oil objective. Images were analyzed in MetaMorph software. Intensity threshold was manually determined for each colocalization marker and kept constant for a binary mask of the marker. The binary mask was overlayed with PD-L1 staining resulting in PD-L1 stain only in pixels positive for the marker proteins. For each cell, integrated intensity of PD-L1 staining in the original images as well as the overlayed images were calculated. Ratio of overlayed intensity over original intensity was used as “fraction under mask”. Unpaired t-test was performed on raw data.

Quantitative RT-PCR:

Cells were lysed after indicated treatments in each experiments and RNA was extracted using RNeasy kit, and cDNA was prepared from extracted RNA using the RNA to cDNA EcoDry Premix. Quantitative RT-PCR was performed using appropriate primer pairs indicated in Supplementary Table 4. 2(-Delta Delta C(T)) method was used to quantify gene expression.

Western blotting:

Cells were lysed in laemmli buffer with protease/phosphatase inhibitors and the lysate was boiled at 95°C for 7 minutes. Lysates were run on 8% or 10% SDS gels and transferred onto nitrocellulose membranes. The membrane was blocked with 5% milk or Licor blocking buffer for 1 hr at room temperature, then incubated with primary antibody overnight at 4°C followed by secondary antibody for 1 hr at room temperature. Blots were visualized using Licor Odyssey Fc Imager for two color or Thermofisher MyECL Imager for single color assays and analyzed on ImageStudio or ImageJ software respectively. Actin or GAPDH expression was used as loading control where indicated. One sample t-test was performed on normalized data for comparison of two groups. One-way ANOVA and post-hoc paired ratio t-test was performed on raw data for three or more groups. For degradation assays, K rate constant for the non-linear regression and slopes for linear regression were used to calculate statistical difference.

Extracellular vesicle, whole cell extract mass spectrometry:

The samples were digested in solution with trypsin overnight at 37C following reduction with 5mM DTT and 14mM alkylation with iodoacetamide. The digests were vacuum centrifuged to near dryness and desalted by Sep-Pak column.

A Thermo Fisher Scientific EASY-nLC 1000 coupled on-line to a Fusion Lumos mass spectrometer (Thermo Fisher Scientific) was used. Buffer A (0.1% FA in water) and buffer B (0.1% FA in ACN) were used as mobile phases for gradient separation (1). A 75 µm x 15 cm chromatography column (ReproSil-Pur C18-AQ, 3 µm, Dr. Maisch GmbH, German) was packed in-house for peptides separation. Peptides were separated with a gradient of 3–28% buffer B over 110 min, 28%-80% B over 10 min at a flow rate of 300 nL/min. The Fusion Lumos mass spectrometer was operated in data dependent mode. Full MS scans were acquired in the Orbitrap mass analyzer over a range of 300-1500 m/z with resolution 120,000 at m/z 200. The top 15 most abundant precursors with charge states between 2 and 5 were selected with an isolation window of 1.4 Thomsons by the quadrupole and fragmented by higher-energy collisional dissociation with normalized collision energy of 35. MS/MS scans were acquired in the Orbitrap mass analyzer with resolution 15,000 at m/z 200. The automatic gain control target value was 1e6 for full scans and 5e4 for MS/MS scans respectively, and the maximum ion injection time is 60 ms for both.

The raw files were processed using the MaxQuant computational proteomics platform version 1.5.5.1 (Max Planck Institute, Munich, Germany) for protein identification. The fragmentation spectra were used to search the UniProt human protein database (downloaded on 01/16/2021). Oxidation of methionine and protein N-terminal acetylation were used as variable modifications for database searching. The precursor and fragment mass tolerances were set to 7 and 20 ppm, respectively. Both peptide and protein identifications were filtered at 1% false discovery rate based on decoy search using a database with the protein sequences reversed.

PD-L1-APEX2 mass spectrometry:

Peptides from streptavidin purified proteins were released via on-bead digestion using trypsin (Promega) and desalted on C18 STAGE Tips [PMID: 12585499]. Eluted peptides were dried and re-suspended in 5% formic acid (FA). Mass Spectrometric analysis was performed on a Thermo Orbitrap Fusion mass spectrometer equipped with an Easy nLC-1000 UHPLC. Peptides were separated with a 115 min gradient of 5–25% acetonitrile in 0.1 % FA at 350 nl/min and introduced into the mass spectrometer by electrospray ionization as they eluted off a self-packed 40 cm, 100 µm (ID) column packed with 1.8 µm, 120 Å pore size, C18 resin (Sepax Technologies, Newark, DE). The column was heated to 60°C. Peptides were detected using a data-dependent method. For each 3 sec cycle, high resolution MS1 scans performed in the orbitrap and the most abundant ions per were isolated, fragmented by CID, and scanned in the ion trap. Ions selected for MS2 analysis were excluded from reanalysis for 60 sec. Ions with +1 or unassigned charge were also excluded from analysis. MS/MS spectra were matched to peptide sequences using COMET (version 2019.01 rev. 5) (PMID: 23148064) and a composite database containing the 20,415 Uniprot reviewed canonical predicted human protein sequences ([http://uniprot.org](http://uniprot.org/), downloaded 5/1/2019) and its reversed complement. Search parameters allowed for two missed cleavages, a mass tolerance of 25 ppm, a static modification of 57.02146 Da (carboxyamidomethylation) on cysteine, and a dynamic modification of 15.99491 Da (oxidation) on methionine. Peptide spectral matches (PSMs) were filtered to 1% FDR using the target-decoy strategy^56^ and then to 1% protein FDR. Label-free quantification was performed using peptide intensities from the integrated areas under each corresponding extracted-ion-chromatogram peak. Intensities for all peptides mapping to each protein were summed for each sample.
